# An Artificial Intelligence Model for Longitudinal Assessment of TCR Repertoires in SARS-CoV-2 Vaccine Recipients

**DOI:** 10.64898/2026.09.16.752071

**Authors:** Zhonghuang Wang, Yu Zhao, Xiao Xiao, Bing He, Yanling Sun, Shan Xiong, Chenchen Qin, Zhipeng Zhou, Lianpeng Chang, Jing Bai, Wenming Zhao, Wenhua Liang, Jianhua Yao

## Abstract

T cells play a crucial role in reducing disease severity during SARS-CoV-2 infection and in shaping long-term immune memory. However, the precise molecular immune responses, particularly involving T-cell receptor (TCR) repertoire changes after full vaccination, and the use of TCR analysis to evaluate vaccine efficacy, remain incompletely understood. In this study, we developed the DeepAir-Cov19 model (AUC=0.94), a large language model tailored to identify SARS-CoV-2-specific TCRs, and observed significant differences in the TCR profiles between antibody-negative and antibody-positive populations before and after vaccination, indicating that the immune status of pre-vaccine recipients can directly assess the efficacy of vaccination. Notably, SARS-CoV-2-specific TCRs expanded to peak levels after the second dose and remained detectable in most subjects up to 10 months post-vaccination. Meanwhile, we also identified three specific V genes, 20 V-J combinations, and 3 epitopes associated with these responses. Finally, by leveraging vaccine-specific TCRs as novel biomarkers, we developed a vaccine efficacy model that predicts antibody levels with a mean AUC of 0.96. These findings underscore the accuracy of the large model in predicting SARS-CoV-2-specific TCRs and reveal a strong correlation between SARS-CoV-2-specific TCR responses and antibody levels. This highlights the complementary and synergistic roles of T cells and antibodies in providing vaccine-mediated protection. Our results offer valuable insights into the longitudinal dynamics of SARS-CoV-2-specific TCRs and illustrate the potential for developing potency evaluation models based on these TCR insights.

## Background

The coronavirus disease 2019 (COVID-19) pandemic, caused by the severe acute respiratory syndrome coronavirus 2 (SARS-CoV-2), has had a devastating impact on global health, with over 7.1 million lives lost(1). SARS-CoV-2 is highly contagious, spreading through respiratory droplets and contact with contaminated surfaces, making it a significant public health concern. COVID-19 symptoms range from mild to severe, with the most common including fever, cough, fatigue, breathing difficulties, and loss of smell. The pandemic has sparked an urgent global effort to develop vaccines targeting SARS-CoV-2, aimed at mitigating its impact on both public health and the global economy. Several novel vaccine platforms have been developed, including messenger RNA (mRNA), viral vectors, protein subunits, and inactivated or attenuated virus vaccines. Given the critical role of the Spike glycoprotein (S protein) in the interaction between the virus and the angiotensin-converting enzyme 2 (ACE2) receptor, as well as the receptor-binding domain (RBD), most researchers have focused on the S protein or RBD as prime targets for vaccine development(2–4). The world has seen an unprecedented number of authorized SARS-COV-2 vaccines, with 50 fully approved for use in several countries(5). Notably, Gao et al.(6) were the first to demonstrate that an inactivated vaccine provided complete protection to non-human primates by inducing effective humoral immune responses. The efficacy and immunogenicity of the three inactivated SARS-COV-2 vaccines currently deployed in China—BBIBP-CorV, CoronaVac, and WIBP-CorV—have been supported by multiple clinical trials(7–11).

As vaccines continue to be distributed worldwide, it remains crucial to measure vaccine response and efficacy to ensure their safety and accessibility(12). While epidemiological data often serve as the gold standard for assessing vaccine effectiveness, the time and effort required to collect and analyze this data make it challenging to quickly adjust and test vaccination strategies. One common method for evaluating vaccine response involves detecting antibodies against viral proteins in serum, typically through ELISA (enzyme-linked immunosorbent assay) or by assessing activated B and T cells(13). However, antibody-based assays have limitations, as detectable antibody levels may take several weeks to develop following vaccination(14). Furthermore, the correlation between serology results and long-term protective immunity remains unclear(15). While neutralizing antibody (nAb) titers can indicate immune protection against SARS-CoV-2 infection(16), assessing them poses challenges due to assay complexity, biohazard risks, and potential issues with durability. ELISpot assays, which measure specificity indirectly, may also yield misleading results if antigen-specific cells remain unactivated(17).

Multiple studies provided compelling evidence that effective prophylactic vaccines against replicating viruses should elicit robust cellular T-cell immunity(18,19). Most vaccines have been shown to induce Th1-skewed T-cell responses, and early T follicular helper (Tfh) and Th1 CD4+ cell responses are associated with the development of effective nAb responses after the initial vaccine dose(20). Additionally, expanded T cell clones detected post-vaccination are predominantly memory cells, contrasting with the effector cells observed during acute natural infection(21). Recent studies have highlighted the critical role of T-cell responses in controlling SARS-CoV-2 infection(22), underscoring the importance of analyzing the T cell receptor (TCR) repertoire to assess vaccine efficacy. T-cell responses can be detected as early as five days post-infection or vaccination, while antibody responses typically take longer—often 20 days or more—to appear. Therefore, TCR immune repertoire testing may offer a more efficient approach than antibody testing. Additionally, T-cell responses tend to have greater longevity than antibody responses, allowing for the evaluation of long-term immune memory formation.

More recently, TCR sequencing (TCR-seq), an application of next-generation sequencing (NGS), has enabled researchers to characterize the diversity of the adaptive immune response using as little as 1–2 mL of whole blood(23,24). The immense variability within the complementarity-determining region 3 (CDR3) of TCR sequences results in most TCRs being unique to each individual. However, a subset of TCRs, known as public TCRs, refer to T-cell receptors that are shared by multiple individuals within a population and are commonly observed in the immune repertoire. These public TCRs, which may arise due to exposure to a shared antigen, have been reproducibly detected in COVID-19 patients and show potential as biomarkers for disease identification and monitoring(25,26). By analyzing immune repertoire changes post-vaccination, researchers can gain valuable insights into vaccine-induced immune responses, including T-cell dynamics. For example, AZD1222 vaccination has been shown to induce a polyfunctional Th1-dominated T cell response, with broad CD4+ and CD8+ T cell coverage across the SARS-CoV-2 spike protein(27). However, distinct changes in clonal composition within the TCR repertoire were observed 28 days after vaccination(28). Another study found that vaccination predominantly caused the expansion of small-sized T-cell clones, leading to a pronounced decrease in clonal diversity compared with natural infection(29).

In this study, we analyzed TCR immune repertoires and antibody responses at four time points over ten monthsin 39 healthy participants who received two doses of an inactivated vaccine. By building a DeepAir-Cov19 deep learning model to identify more SARS-CoV-2 specific CDR3 sequences, we found significant differences in TCR between antigen-negative and antigen-positive populations before and after vaccination, indicating that the immune status of pre-vaccine recipients can directly assess the efficacy of vaccination. A significant amplification of the TCR response after the second dose of vaccine was also observed, achieving a faster response compared to the 28-day timeline for antibody production-the TCR response was detectable within only 7 days. We not only demonstrated the feasibility of using TCR to assess vaccine response, but also constructed machine learning models to provide TCR-based alternatives for nAb detection. These findings indicate that immune repertoire sequencing has the potential to be a powerful tool for the assessment of vaccine efficacy, particularly for the assessment of T-cell responses. By providing a comprehensive view of the immune system’s response to vaccination, immune repertoire sequencing can aid in the development and optimization of vaccines against infectious diseases such as COVID-19 and provide a rapid and robust approach to vaccine evaluation.

## Methods

### Materials preparation

#### Study participants and vaccinations

Forty-two participants were enrolled in this study, which investigated the efficacy of an inactivated SARS-CoV-2 vaccine funded by Beijing Geneplus Technology Co Ltd. The trial specifically recruited healthy adults who had not been previously infected with SARS-CoV-2. The SARS-CoV-2 inactivated vaccine utilized in the trial, primarily comprising Beijing Kexing Zhongwei (Vero cells) and Beijing Biological (Vero cells) (Supplementary Tables S1), was administered intramuscularly in the participants’ arms. Throughout the trial, the health conditions and any adverse events experienced by the participants were self-reported and closely monitored by the investigator. The participant group consisted of males and females, with ages ranging between 25 and 40. Detailed demographic characteristics of the participants can be found in Supplementary Table S1. All participants provided written informed consent, and the study was approved by the Ethics Committee of Guangzhou Laboratory Medical Research and Lunar Review 2021, No. 78

#### PBMC collection

Peripheral blood mononuclear cells (PBMCs) were isolated from whole blood collected in vacutainer tubes. Blood samples were mixed with an equal volume of PBS buffer to prevent cell clumping. The samples were then transferred to 15 mL lymphocyte separation tubes. If the separation solution stained the tube walls, the tubes were centrifuged at 800×g for 1 minute at 20°C to ensure the solution settled. Samples containing cell clots were excluded. For the first centrifugation, samples were centrifuged at 800×g for 15 minutes at 20°C. The upper plasma and PBS layers were removed, and the PBMC layer was carefully transferred to a new centrifuge tube. For the second centrifugation, samples were centrifuged at 16,000×g for 2 minutes at 20°C. After removing the supernatant, 1 mL of RNAlater reagent was added to the PBMCs. The cells were pipetted until fully dispersed, yielding PBMCs ready for further processing.

#### Chemiluminescent Magnetic Bead Immunoassay(neutralizing antibodies against SARS-CoV-2)

The antibody detection method uses a competition method in which the neutralizing antibody in the sample competes with the biotinylated 2019-nCoV-specific antibody for the acridinium ester-labeled S protein to form a “biotinylated 2019-nCoV-specific antibody-acridinyl ester S protein” complex, and streptavidin-coated magnetic particles are added to form a “biotinylated 2019-nCoV-specific antibody-acridinyl ester S protein” complex through the interaction between biotin and streptavidin. After washing and removal of substances that do not bind to the magnetic particles, pre-excitation solution and excitation solution are added to the reaction mixture. The resulting chemiluminescence signal is measured and expressed in relative luminescence units (RLU). The concentration of the neutralizing antibody in the sample is inversely proportional to the RLU detected by the instrument. The concentration of neutralizing antibodies in the sample is also calculated from the standard curve.

When loading the kit onto the luminometer for the first time, the reagents need to be mixed to re-suspend any microparticles that may have settled during shipment. The calibrator should be equilibrated to room temperature after removal from 2∼8°C and gently turned over and mixed before use; after use, cap the bottle tightly and return the calibrator to storage at 2∼8°C. Execute the calibration command: Execute the calibration command message, if the calibration is valid after calibration, continue to execute the subsequent procedures; if the calibration is invalid after calibration, re-execute the calibration command message. Check the sample volume in the sample cup to ensure that the sample volume in the sample cup is kept above 100uL before each test. Finally, send the sample into the luminescent instrument for testing.

#### High-throughput sequencing of TCR β-chain and identification

Sequencing of the CDR3 regions of human T-cell receptor beta (TCR-β) chains was conducted using a multiplex PCR amplification protocol. Two sets of PCRs, namely PCR1 and PCR2, were performed on DNA extracted from collected peripheral blood mononuclear cells (PBMC). The initial PCR employed a mixture of multiplexed V- and J-gene primers to amplify all potential recombined receptor sequences from the DNA sample. The PCR conditions were as follows: an initial denaturation cycle at 95°C for 15 minutes, followed by 10 cycles of denaturation at 94°C for 30 seconds, annealing at 60°C for 90 seconds, and extension at 72°C for 30 seconds. Subsequently, a second PCR was conducted using universal primers. Afterward, the PCR products were subjected to sequencing using an Illumina MiSeq platform, employing a 150-cycle paired-end protocol and sequence-ready primers. The resulting raw data was transferred to Adaptive Biotechnologies for further processing. A comprehensive report was generated, encompassing samples that passed quality checks, along with a normalized and annotated TCR-β profile repertoire. The raw reads underwent subsequent processing and analysis using the following steps: (1) removal of sequencing reads lacking the primers for multiplex PCR using Cutadapt(55); (2) merging the remaining high-quality paired-end reads to generate contigs using Pear(56); and (3) identification of the CDR3 region using MiXCR(57) with default parameters. The CDR3 sequence was defined based on the ImMunoGeneTics (IMGT) V, D, and J gene references, with the CDR3 region encompassing the amino acids between the second cysteine of the V region and the conserved phenylalanine of the J region. A comprehensive report including samples that passed quality checks, along with a normalized and annotated TCR-β profile repertoire was generated.

### Machine learning model for predicting antibody levels based on TCR data

#### Identification of antibody-associated CDR3 sequences of TCRs

We have devised a framework that incorporates statistical learning and machine learning techniques to identify T-cell receptor sequences (TCRβs) associated with specific subject phenotypes.

To evaluate the significance of the association between each feature (TCRβ) and the class assignment (phenotype status) in the training data, we employ a T-test. This involves calculating the p-value by comparing the frequencies of the TCRβ in question across different classes. By analyzing the frequency distributions of TCRβs, we identify a collection of phenotype-associated TCRβs that comprise both significant antibody-positive TCRs and antibody-negative TCRs. Although FDR could not be performed due to the sparsity of the matrix, the use of supervised learning for feature selection proved to be effective. In particular, the lasso regression approach was found to be more suitable for the feature selection of sparse matrices than decision tree models such as RF, GDBT, and XGBoost.

Lasso regression is a linear model that uses this cost function:

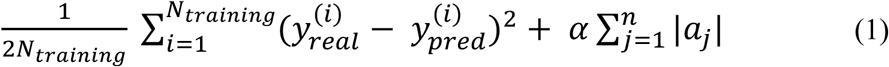

The coefficient *a*_j_ represents the weight assigned to the j feature. The final term L_1_ penalty, also known as the regularization term, is denoted by α, which is a hyperparameter used to control the strength of this penalty. The cost function increases as the coefficients of the features become larger. Therefore, the objective of Lasso regression is to optimize the cost function by minimizing the absolute values of the coefficients. The appropriate value of the α hyperparameter is determined through cross-validation. By minimizing the cost function, Lasso regression automatically selects the most relevant features. To perform feature selection, we apply Lasso regression to a scaled version of the dataset and retain only those features with nonzero coefficients. It is important to fine-tune the α hyperparameter to ensure the effectiveness of the Lasso regression. In our model, the α value is determined to be 6.997 × 10⁻⁶. Following the feature selection process, out of the original 1,173,221 CDR3s in the training set, only approximately one hundred CDR3s remained for further analysis. Finally, a total of 528 CDR3 features were obtained after ten repeated experiments.

#### Machine learning method for classification of antibody levels

Comparing CNN model, TCRdist and GLIPH2 method, the logistic regression shows better performance. For TCR repertoires classification, logistic regression was found to outperform other machine learning models such as RF and SVM methods in combination with two encodings (3-mer and 4-mer encoding).

Logistic regression is a supervised machine learning algorithm primarily utilized for classification tasks, aiming to predict the probability of an instance belonging to a specific class. It is commonly employed in classification algorithms and is known as logistic regression. Despite its name, it is categorized as regression due to its utilization of the linear regression function’s output as input and the application of a sigmoid function to estimate the probability for the given class. Unlike fitting a straight line or hyperplane, the logistic regression model employs the logistic function to confine the output of a linear equation within the range of 0 to 1. The sigmoid function is defined as:

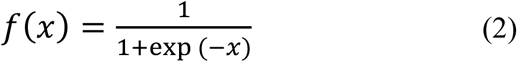

The Logit function maps y to the sigmoid function of x.

### Identification of SARS-Cov-2 Specific CDR3 clones

#### DeepAir-Cov19

DeepAir-Cov19 is an artificial intelligence model designed to identify SARS-Cov-2-specific CDR3 sequences based on DeepAIR(33). It comprises three feature encoders that play a crucial role in capturing essential information from the input data. Additionally, DeepAir-Cov19 incorporates a gating-based attention mechanism, which helps identify and extract significant features. To further enhance its predictive capabilities, a tensor fusion mechanism is employed to effectively integrate the extracted features. By leveraging these advanced techniques, DeepAir-Cov19 demonstrates its ability to accurately predict the binding specificity between TCR and SRAS-Cov-2 antigens. We downloaded experimentally validated data of CDR3 binding associations from natural and synthetic exposure to SARS-CoV-2 from the Immune Access project(32). These CDR3 were used as positive training cases, and the CDR3 that were selected from the unvaccinated healthy population without positive cases were used as negative cases. During the training phase, DeepAir-Cov19 employs two loss functions to optimize its performance. The main loss function is the mean squared error (MSE) loss, denoted as ℒMSE, which is utilized by the main regression layer. This loss function encourages DeepAir-Cov19 to directly predict an accurate specificity score for binding. Additionally, an auxiliary loss function called categorical cross-entropy (CE) loss, denoted as ℒCE, is employed by the auxiliary specificity grading layer. The purpose of this loss function is to facilitate DeepAir-Cov19 in learning the accuracy of specificity orders. The total loss function for the prediction of binding specificity in DeepAir-Cov19 is defined as follows:

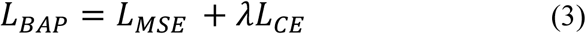

Finally, the DeepAir-Cov19 model is able to predict unseen CDR3 sequences and discriminate whether the sequence has a specificity to the SARS-CoV-2 epitopes. If the prediction value is larger than 0.5, then the sequence has a specificity with the SARS-CoV-2 epitopes and the sequence is considered to be a SARS-CoV-2-associated CDR3. In this study, in order to improve the discriminant accuracy, we increased the threshold to 0.8.

#### Unsupervised Clustering method for CDR3 sequences tracing

Fuzzy c-means algorithm (FCMA) clustering analysis is an important clustering method for unsupervised machine learning. In contrast with hard clustering, soft clustering methods can assign a gene to several clusters which can overcome the shortcomings of conventional hard clustering techniques and offer further advantages. The R package of Mfuzz(34) was used to calculate the clustering for the analysis of the temporal CDR3 changes.

All DeepAir-Cov19-TCR CDR3 sequences are clustered in different groups with different fate developments. In the clustering, the group with the highest number of sequences was considered to be the most consistent with the trend of CDR3 fate development after vaccination.

## Results

### Study Design and Sustained Immune Protection in Females

To comprehensively assess the humoral immune response to SARS-CoV-2 vaccination, we enrolled 42 healthy participants who received two doses of inactivated vaccines—either the Beijing Kexing Zhongwei (Vero cells) or the Beijing Bio (Vero cells). These participants were part of two studies (Figure 1A). The first study focused on the long-term analysis of SARS-CoV-2-specific TCRs, as predicted by DeepAir-Cov19. TCR-seq was conducted on 39 participants, yielding a total of 156 blood samples collected at four time points: before vaccination (T0; n = 39), two weeks after the first dose (T1; n = 39), one week after the second dose (T2; n = 39), and ten months following the initial vaccination (T3; n = 39). The second study aimed to predict antibody levels based on TCR repertoires ten months after the initial doses, including 40 participants. Notably, 37 participants participated in both studies, with four rounds of TCR-seq and antibody assays conducted at T3.

**Figure 1.**
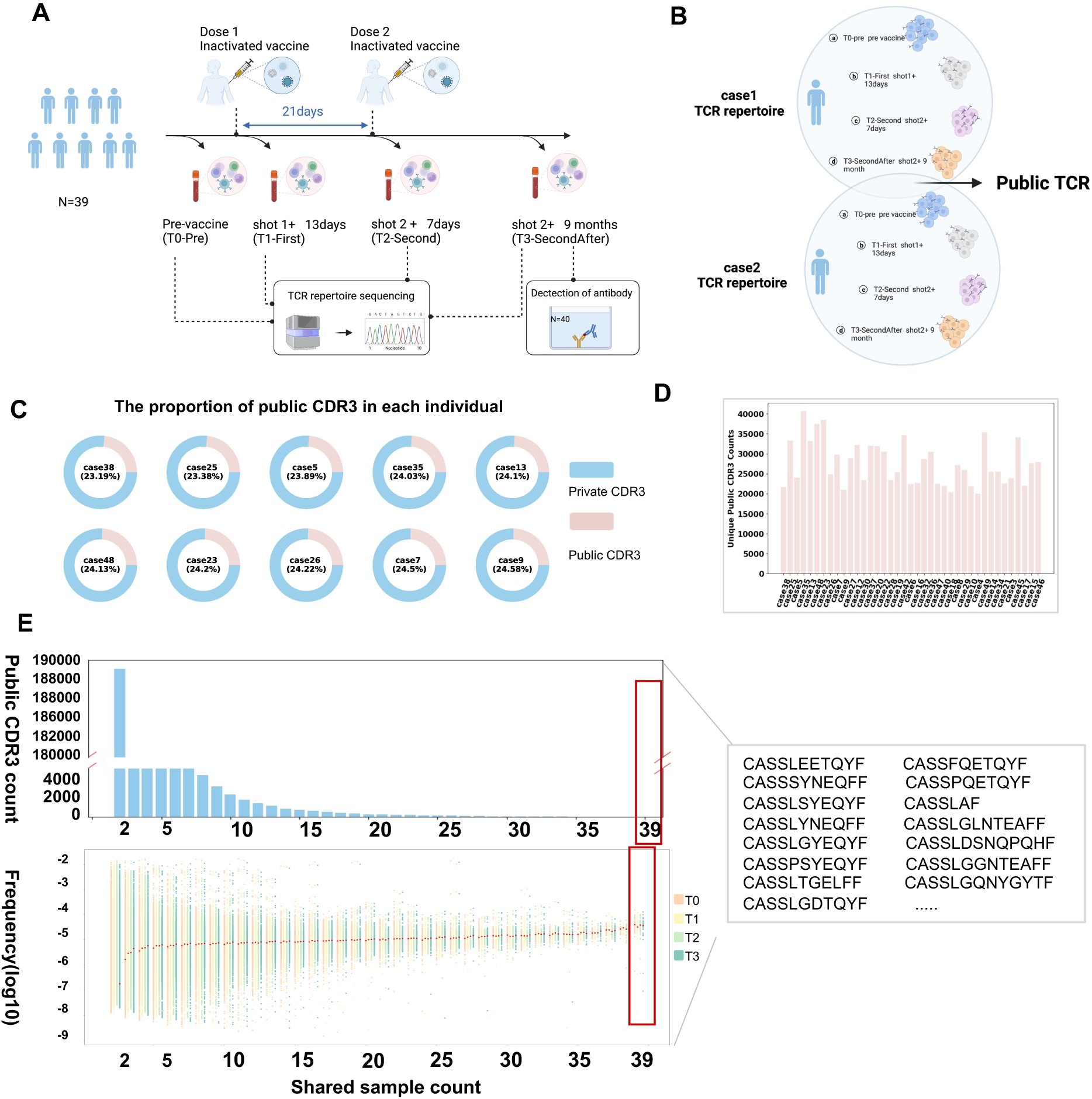
Study design and the feature of public TCR CDR3 sequences. (**A**) This study included 42 participants, of whom 39 had complete TCR repertoire records at four time points, and 40 had antibody level records. Participants received two doses of inactivated vaccines with a 21-day interval. PBMCs were collected from included subjects at the following four-time points: Pre vaccination (Pre -T0), 13 days after the First dose vaccination (denoted as First-T1), 7 days after the Second dose vaccination (Second-T2), and 9 months after the Second dose (SecondAfter-T3). Peripheral blood mononuclear cells (PBMCs) were collected at four time points. PBMCs were analyzed via TCR immune repertoire sequencing using NGS and antibody assays. Antibody levels were assessed at the final time point. (**B**) Combined TCR Immune Repertoires and Public CDR3 Definition. The combined TCR immune repertoire includes all TCRs of an individual at four different times. Public CDR3 sequences are defined as those shared by more than one individual. (**C**) Proportion of Public CDR3 Sequences. Pink represents public CDR3 sequences. Blue represents private CDR3 sequences (**D**) Counts of unique public CDR3 sequences (pink) in each participant’s repertoire. (**E**) Distribution of Public CDR3 Sharing Levels. The number of public CDR3 sequences shared among different numbers of participants. The mean frequencies of amino acid (aa) sequences in the CDR3 region were plotted against their sharing levels, with each dot representing the average frequency of a specific aa sequence among participants. The graph shows four time points (T0: orange, T1: yellow, T2: green, T3: blue), with red dots indicating the median value for each sharing category.

The demographic characteristics of the participants are summarized in Table 1. Participants were between 25 and 40 years old, and age did not significantly correlate with antibody response (p = 0.15). However, a notable gender difference was observed: female participants exhibited significantly higher antibody levels than male participants at T3 (Fisher’s exact test, p = 0.002), consistent with findings from previous studies(30). Specifically, all male participants had negative antibody assay results, while more than half of the female participants had positive results (Supplementary Table 1). These findings suggest that inactivated vaccines may elicit a stronger immune response and more persistent protection in females. Notably, we chose to analyze T cell receptor sequencing at a sampling time of 1 week after the second dose of vaccine (T2), which is different from previous studies that confirmed antibody production takes 3 weeks after the second dose. Our final sampling was conducted after a ten-month follow-up of the initial vaccine doses, a longer observation period than the six month follow-up in the previous study(31).

**Table 1.** Demographic characteristics of study participants.

| **Inactivated-vaccinated donors for four times** |  |  |
| --- | --- | --- |
| **Characteristic** |  | **Total(N = 39)** |
| Median age – year (range) |  | 30(25-40) |
| Sex – no. (%) | Female | 27(69.2) |
|  | Male | 12(30.8) |
| Median time between vaccine doses – days |  | 21(21-22) |
| Median time between before first dose and sample collection – days |  | 1 |
| Median time between first dose and sample collection – days |  | 13 |
| Median time between second dose and first sample collection – days |  | 7 |
| Median time between second dose and second sample collection – days |  | 274 |
| **Inactivated-vaccinated donors for detection of antibody** |  |  |
| **Characteristic** |  | **Total(N = 40)** |
| Median age – year (range) |  | 30(25-40) |
| Sex – no. (%) | Female | 27(69.2) |
|  | Male | 12(30.8) |
| Antibody - no. (%) | Positive | 17(42.5) |
|  | Negative | 23(57.5) |
| Median time between vaccine doses – days |  | 21(21-22) |
| Median time between before first dose and sample collection – days |  | 1 |
| Median time between first dose and sample collection – days |  | 13 |
| Median time between second dose and first sample collection – days |  | 7 |
| Median time between second dose and second sample collection – days |  | 274 |

### The Feature of Public TCR CDR3 sequences

We analyzed the collective immune repertoires of 39 participants, which included all TCR sequences at four time points. A total of 5,158,389 unique CDR3 sequences were identified across these repertoires, of which 32,141 unique public sequences were shared by more than two participants, (Figure 1B). Public CDR3 sequences represented approximately 20% of each participant’s repertoire (Figure 1C), with counts ranging between 20,000 and 40,000 (Figure 1D, Supplementary Figure 1). The remaining 90.85% of sequences were private CDR sequences, found in only one participant. Among the public sequences, 189,107 were shared by two participants, and 34 were shared by all 39 participants (Figure 1E). The degree of sharing ranged from private to highly public, with the frequency of each CDR3 sequence correlated positively with the level of sharing. Sequences shared among more participants tended to be more abundant, as reflected in the increasing median frequency observed with higher levels of sharing (Figure 1E). These public CDR3 sequences play a pivotal role in understanding the TCR immune repertoire response following vaccination.

### DeepAir-Cov19: A Deep Learning Model for SARS-CoV-2-Specific TCR Recognition

Despite the vast diversity of existing TCRs, experimentally validated SARS-CoV-2-specific TCRs remain scarce. To address this gap and further investigate the TCR immune repertoire to vaccination, we developed a deep learning model DeepAir-Cov19 designed to identify novel TCR sequences with specificity to SARS-CoV-2 antigens, with the potential to uncover key TCRs associated with vaccine responses.

For the model data (Table 2), we use publicly available datasets containing a total of 250,204 TCR clones. As positive examples, we incorporated experimentally validated SARS-CoV-2-specific TCR sequences from the SARS-CoV-2-MIRA dataset(32), which contains 125,102 TCR clones specific to SARS-CoV-2 antigens. As negative examples, we used TCR sequences from healthy individuals, ensuring that SARS-CoV-2 antigen-specific TCRs were excluded containing 125,102 TCR clones. The dataset was balanced with an equal ratio (1:1) of positive and negative samples to develop a predictive model for SARS-CoV-2 antigen specificity. For model training and evaluation, the data split into three subsets: 70% for training, 20% for validation, and 10% for testing. This split was designed to maintain balanced representation of TCRs binding to SARS-CoV-2 antigens across all sets, ensuring robust model performance.

**Table 2.**
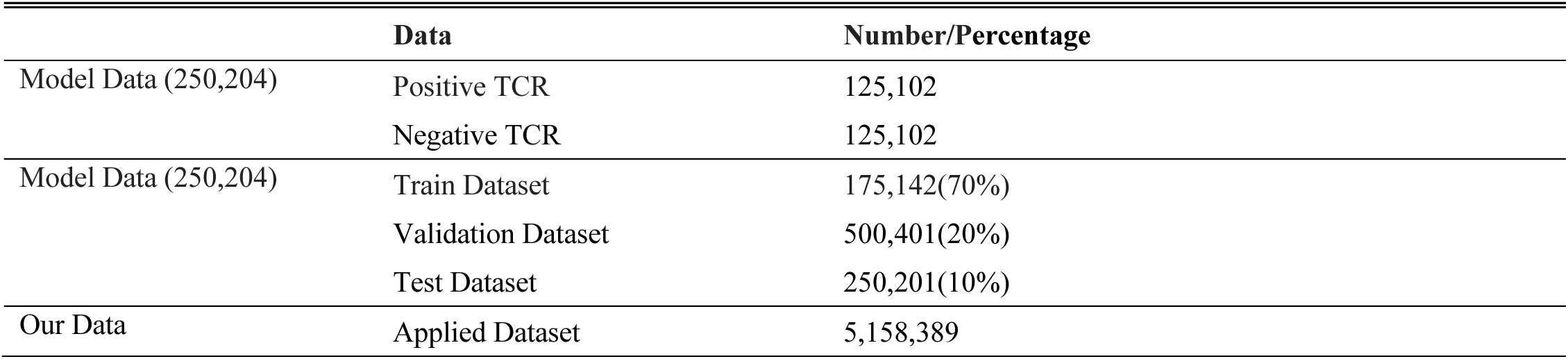
The Model Dataset.

|  | **Data** | **Number/Percentage** |
| --- | --- | --- |
| Model Data (250,204) | Positive TCR | 125,102 |
|  | Negative TCR | 125,102 |
| Model Data (250,204) | Train Dataset | 175,142(70%) |
|  | Validation Dataset | 500,401(20%) |
|  | Test Dataset | 250,201(10%) |
| Our Data | Applied Dataset | 5,158,389 |

DeepAir-Cov19 model is a deep learning model for SARS-CoV-2-Specific TCR recognition. For the modeling, we use the SARS-CoV-2 antigen specificity dataset for retraining, based on the DeepAIR(33), making the model applicable to SARS-CoV-2-specific. The DeepAir-Cov19 model consists of three main processing stages: feature embedding, MLP (Multilayer perceptron) head, and task-specific prediction (Figure 2A). During the feature embedding stage, DeepAir-Cov19 employs two feature encoders, namely the gene encoder and sequence encoder, to comprehensively encode the TCR. The gene encoder incorporates the V(D)J gene usage information using a trainable embedding layer. The sequence encoder utilizes a multi-layer transformer to generate a high-level representation of sequence information for the paired chains. Moving to the multilayer perceptron head stage, a module is employed to extract crucial features from the obtained sequence and gene embeddings. For SARS-CoV-2 antigen-specific predictions, the classification layer is employed to perform the prediction task. Additionally, the V(D)J gene usage module is available as an optional feature.

**Figure 2.**
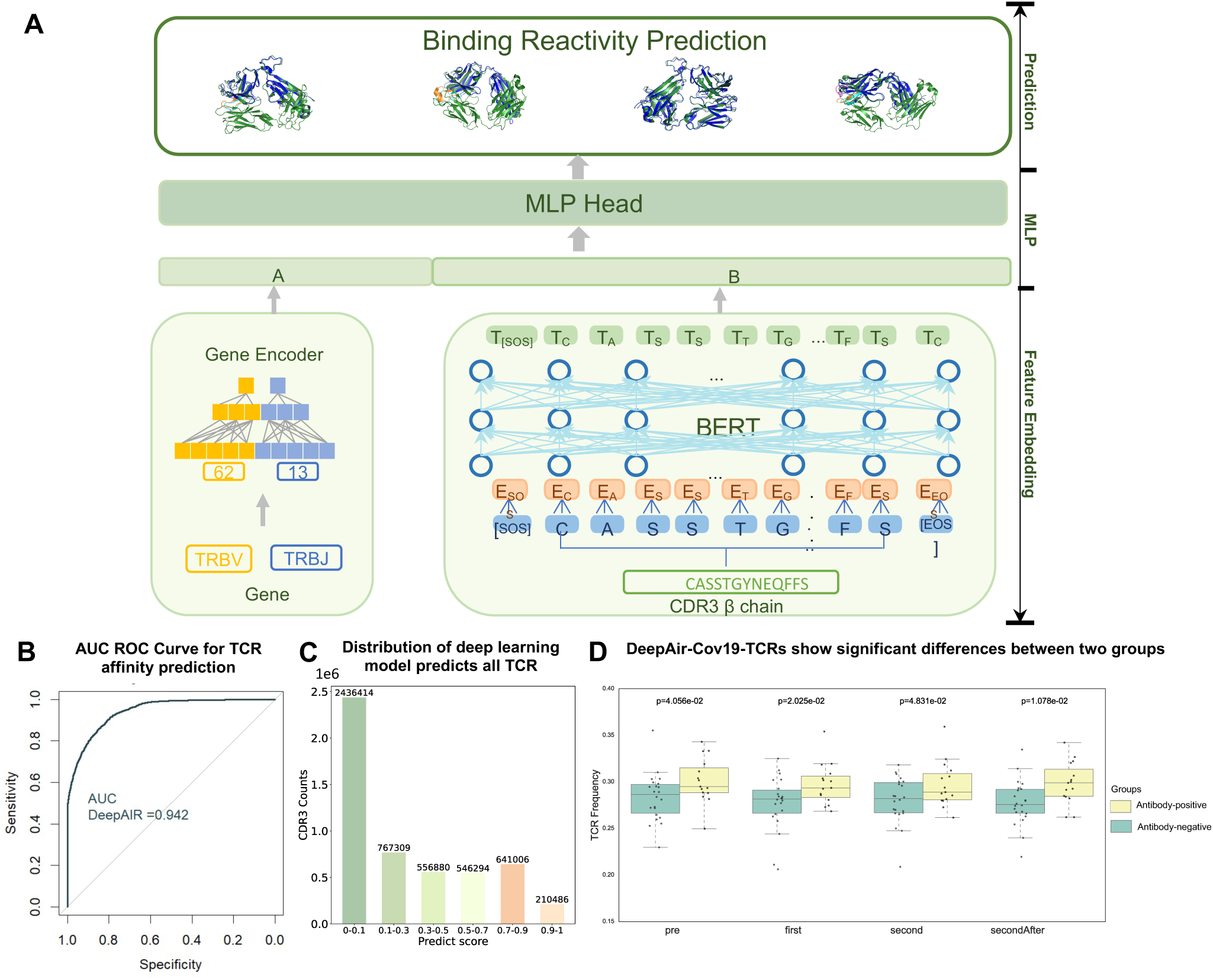
The framework of DeepAir-Cov19 and the feature of DeepAir-Cov19-TCRs. (**A**) The architecture of the DeepAir-Cov19 artificial intelligence model designed to identify SARS-Cov-2-specific CDR3 sequences. The model consists of feature embedding, a multilayer perceptron (MLP) head, and task-specific prediction. Feature embedding extracts information from gene and sequence inputs, which are processed through the MLP to produce representations. The final prediction layers map these representations to identify SARS-Cov-2-specific CDR3 sequences. (**B**) Model performance. ROC curve showing the model’s performance in predicting SARS-CoV-2-specific CDR3s, with an AUC of 0.942. (**C**) Distribution of DeepAir-Cov19 predicted scores for CDR3 sequences in 5,158,389 TCR clones, ranging from 0 to 1. (**D**) Differences between DeepAir-Cov19-TCR in antibody detection negative and positive groups. Between-group differences in total frequency values of DeepAir-Cov19-TCR CDR3 sequences at four time points (T0, T1, T2, T3) between antibody-negative (green) and positive (yellow) participants. Each gray dot represents one participant CDR3 frequency.

Remarkably, the DeepAir-Cov19 model achieved an area under the curve (AUC) of 0.94 for the prediction of SARS-CoV-2 antigen specificity (Figure 2B). Subsequently, we applied the model to our dataset, which comprised 5,158,389 TCR clones derived from 156 TCR immune repertoires, surpassing the quantity of experimentally validated data. Upon evaluating the predicted scores for all TCRs, it was observed that the majority of CDR3 sequences (70.2%) exhibited prediction scores below 0.5, suggesting a low specificity for the SARS-CoV-2 antigen (Figure 2C). To enhance the accuracy of identifying SARS-CoV-2-specific TCR sequences, we defined TCRs with prediction scores above 0.8 as DeepAir-Cov19-TCRs. These high-confidence TCRs were then further analyzed for their potential relationship with antibody detection results. To investigate this, we calculated the summed frequency of DeepAir-Cov19-TCRs for each participant at four time points (Figure 2D). While the antibody detection was conducted at time point T3, we observed significant differences (p < 0.05) in TCR frequencies at earlier time points, even before vaccine administration, in both antibody-negative and antibody-positive groups. Notably, the TCR frequency in the antibody-positive group was consistently higher than in the negative group. These findings suggest that the AI model not only accurately predicts SARS-CoV-2-specific TCRs but also reflects underlying immune responses, which could potentially be used to predict antibody detection outcomes. This may indicate that the immune status of the person before the vaccine can be used to directly evaluate the effect of the vaccine.

### Longitudinal follow-up of two doses: the feature of DeepAir-Cov19-TCRs CDR3 sequences and Gene Usages

Previous studies have focused predominantly on the role of antibodies or nAb responses following vaccination, with little research on the longitudinal development of TCR repertoires. To investigate the temporal dynamics of SARS-CoV-2-specific TCRs, we performed unsupervised clustering of the DeepAir-Cov19-TCRs CDR3 sequences across the four time points using Mfuzz(34) (Figure 3A). Within these clusters, SARS-CoV-2-specific CDR3 sequences were grouped into four distinct clusters. Cluster 1, which contained the highest number of sequences, was deemed most representative of the post-vaccination trend for sequences specific to SARS-CoV-2. Notably, we observed that the number of sequences in Cluster 1 expanded following the administration of the second dose of the vaccine and remained for ten months, surpassing the levels observed after the first dose. This expansion pattern aligns with previous reports on the development of antigen-specific antibody responses post-vaccination.

**Figure 3.**
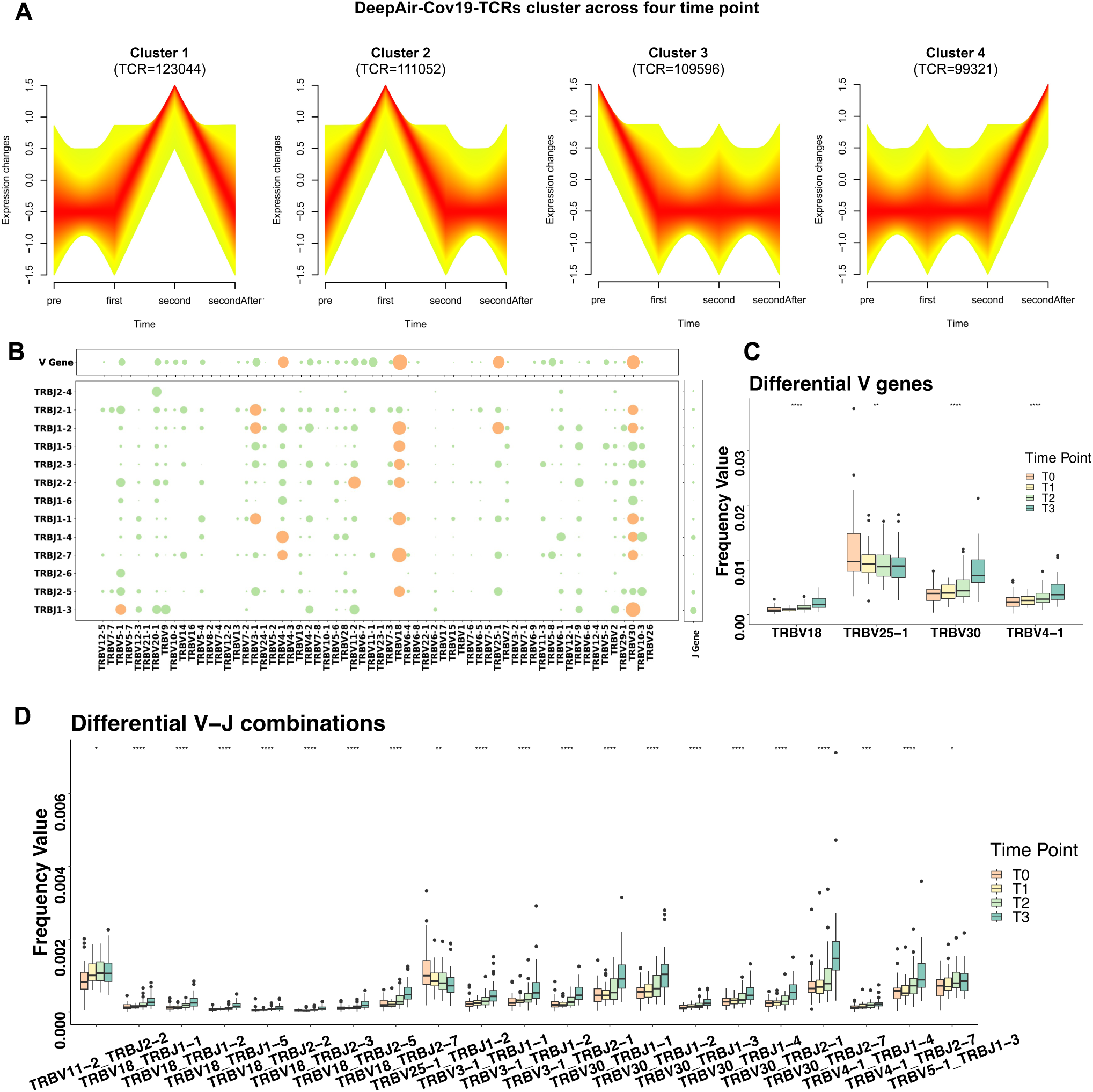
Characteristics of the identified vaccine-specific V genes, J genes, V-J combinations, and corresponding epitopes. (**A**) Longitudinal frequency patterns of DeepAir-Cov19-TCRs CDR3 sequences across four time points (T0, T1, T2, T3). Frequency values were standardized via z-score normalization. The Fuzzy C-Means algorithm was used to perform the clustering. Cluster cores (membership > 0.90) are highlighted in red. (**B**) One-way ANOVA test for differences in frequency of all V genes, J genes, and V-J combinations for the pre-vaccine, post-first shot, and post-second shot periods. The orange dots show significant differences (p<0.05). The green dots represent non-significant differences (p>0.05). (**C**) Differential V genes in three groups (T0 green, T1 yellow, T2 red). There are 4 V differential genes (p<0.05) from 62 V genes, (one-way ANOVA test). (**D**) Differential V-J (Variable-Joining) combinations in three groups (T0 green, T1 yellow, T2 red). There are 21 V-J combinations of differential genes (p<0.05) from 806 V genes (one-way ANOVA test).

V (Variable) and J (Joining) genes are critical components of TCRs and play a key role in the diversity of TCRs, enabling the immune system to recognize a wide range of antigens. In our study, we identified 62 V genes, 13 J genes, and 806 V-J combinations across the first three time points of the study. By analyzing differential gene frequencies using ANOVA, we observed significant differences (p < 0.05) in the expression of some V genes and V-J combinations, while J genes did not exhibit significant changes over time (p > 0.05) (Figure 3B). Among the differentially expressed V genes, four (TRBV18, TRBV25-1, TRBV30, and TRBV4-1) showed significant variations in frequency between the pre-vaccination, post-first dose, and post-second dose periods (p < 0.05) (Figure 3C). Similarly, 21 V-J combinations displayed notable changes in frequency levels (Figure 3D). Notably, while the frequency of TRBV25-1 and its V-J combination TRBV25-1_TRBJ1-2 significantly decreased after vaccination, the rest showed a marked increase. These results suggest that vaccination may selectively influence specific V genes and V-J combinations.

Further analysis revealed that increases in the frequency of 3 V genes and 20 V-J combinations corresponded with a decrease in other combinations, indicating selective expansion of certain TCR clones in response to SARS-CoV-2 antigens. Interestingly, some individuals showed significant amplification of specific V genes and V-J combinations after the second dose (Figure 4A and 4B, Supplementary Figure 2 and 3), suggesting variability in vaccine response between individuals. These findings have important implications for vaccine development and personalized medicine, as they underscore the individual variability in immune responses. Overall, our study provides insights into the molecular mechanisms underlying the development of TCR repertoires in response to the COVID-19 vaccine, highlighting the role of certain V genes and V-J combinations in driving immune responses and the potential for personalized approaches to enhance vaccine efficacy.

**Figure 4.**
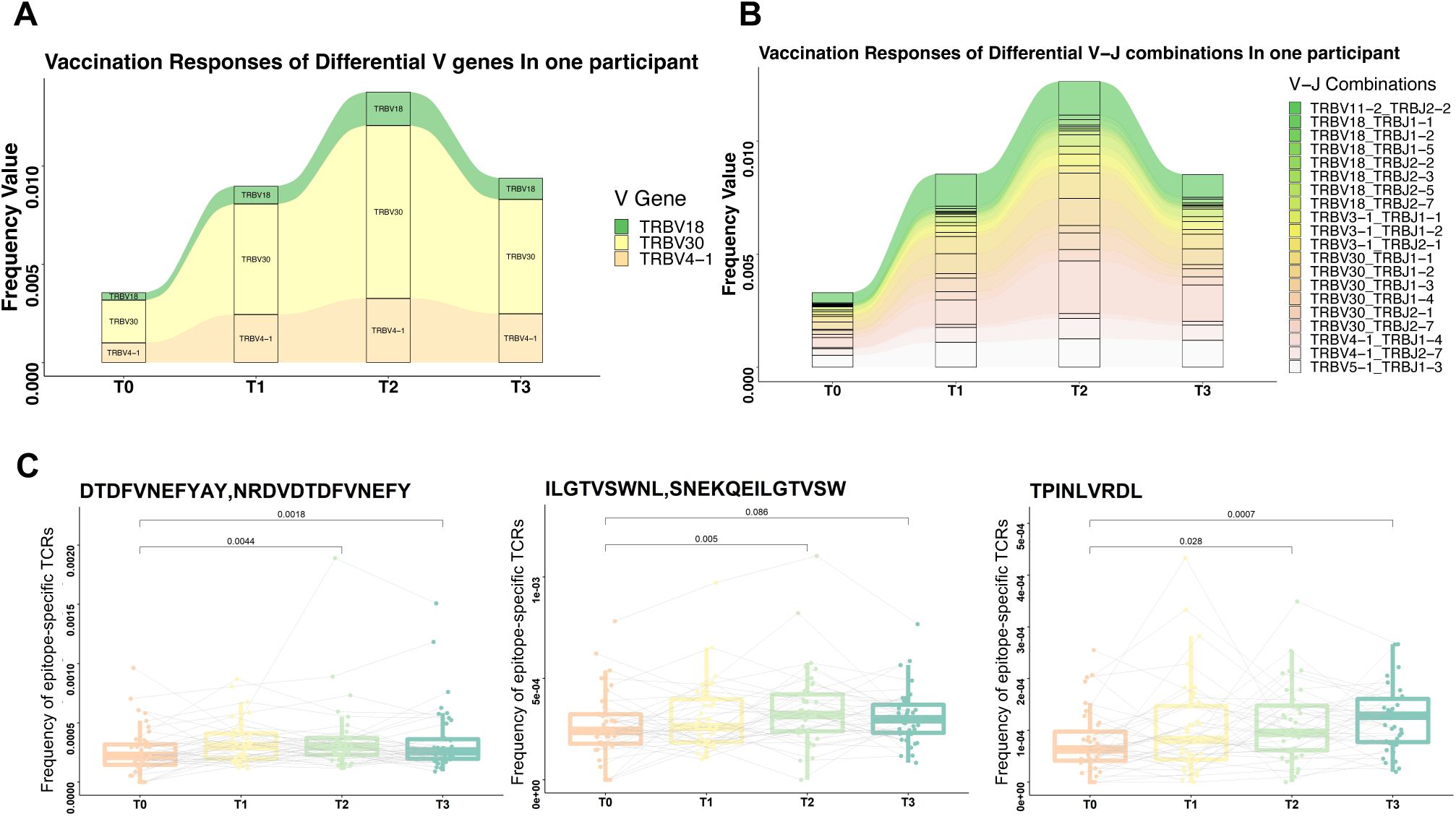
Tracking vaccine-specific V genes, J genes, V-J combinations, and corresponding epitopes. (**A**) Differential V segment genes are shown in vertical stacks with curved segments (thickness proportional to the frequency of TCRs) in one participant. (**B**) Differential V-J combinations genes tracking in one participant for ten months following the initial dose of inactivated vaccine. (**C**) Three epitopes expanded significantly (p<0.05) after two vaccine stimulations. The total frequency of clones, which are binding to a certain epitope, expanded significantly between T0 (pre vaccinations) and T2 (post two doses of vaccinations). Dots represent the total frequency of clones binding to a certain epitope in one participant. A solid gray line represents the development of total clone frequency for one participant. The vertical coordinate shows the sum of clone frequency binding to a certain epitope.

In addition to exploring the longitudinal expansion of TCR from the perspective of CDR3 sequences and V-J combinations, we also considered TCR clones, which combine CDR3 sequences with V-J combinations and exhibit greater diversity. Using the SARS-CoV-2-MIRA dataset containing 125,102 experimentally validated TCR clones with SARS-CoV-2 specificity and corresponding epitope information, we mapped these clones onto the 39 participants’ TCR repertoires. This allowed us to aggregate clones with distinct epitopes within each repertoire. Our analysis identified three epitopes (NRDTDFDFVNEFYAY, SNEKQEILGTVSWNL, and TPINLVRDL) that showed significant expansion (t-test, p < 0.05) seven days after the second vaccination (T2) (Figure 4C). Of these, NRDTDFDFVNEFYAY and SNEKQEILGTVSWNL are located within ORF1ab, while TPINLVRDL is located on the surface protein. These epitopes play crucial roles in the immune response following vaccination, and their significant expansion suggests that the vaccine effectively triggered TCR responses specific to these SARS-CoV-2 epitopes. The identification of TCRs targeting specific epitopes that expanded significantly after vaccination is a key finding, indicating the vaccine’s ability to induce an antigen-specific immune response.

### Machine learning model for identifying antibody-associated CDR3 sequences and predicting classification results

In Section 3, we demonstrated that Deep-Air-Cov19-TCR effectively differentiates between antibody-negative and antibody-positive groups. Building on this, we sought to leverage TCR data to predict antibody status in both groups. Previous studies have highlighted the critical role of antigen-specific TCRs in various diseases, including cancer and autoimmune disorders(35). Although there are studies that have explored the relationship between antibody responses and the CDR3 sequence of the TCR(36), there are limited studies on the TCR to predict antibody status. Given the pivotal role of T cells in the immune response to vaccines, the ongoing COVID-19 pandemic has heightened the need to better understand T cell responses to vaccination. To address this gap, we developed an integrated pipeline for identifying CDR3 sequences that correlate with SARS-CoV-2 antibody detection results. We then applied a machine learning model to classify antibody status based on these characteristic CDR3 sequences (Figure 5A). This approach provides an effective framework for screening and classifying CDR3 sequences in small TCR immune repertoires.

**Figure 5.**
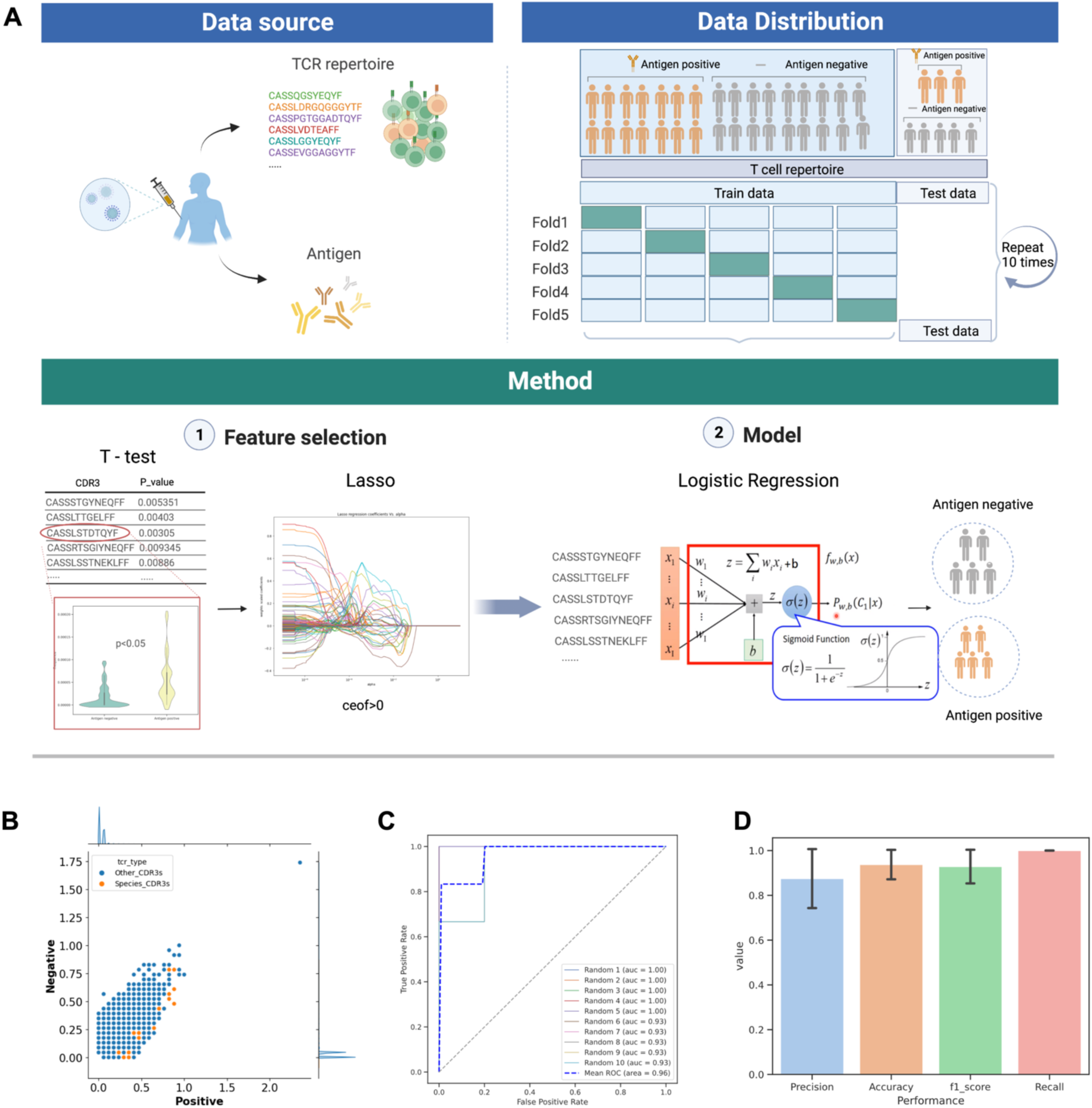
Antibody prediction model performance using vaccine-specific CDR3 sequences. (**A**) Data source: TCR immune repertoires were acquired using small-volume blood samples from 40 individuals after two doses of inactivated vaccines during a ten-month period. Meanwhile, serologic testing was used as ground truth. Data Distribution: 40 participants were divided as the training and test dataset at an 8:2 ratio with a balanced positive-negative distribution. The five-fold cross-validations were conducted ten times by randomly splitting the samples in the training dataset at each time. Method: The feature selection module includes the selection of CDR3 sequences with differences between groups using t-test (p<0.05), and lasso regression to do further feature screening. Classification module: The classification module uses logistic regression for binary classification of antibody assay using CDR3 sequences. (**B**) Distribution of TCRβ CDR3 sequences in antibody positive and negative samples. The size of the spot in each position represents the abundance of CDR3 sequences. Antibody-positive CDR3 sequences identified in this study are represented by orange dots. All other CDR3 sequences identified are shown in gray. (**C**) ROC curves show the classification performance by five-fold cross-validation. The blue line is the mean ROC of the 10 randomized experiments. (**D**) The model’s overall performance is evaluated using key metrics, including precision, recall, F1 score, and accuracy.

In our study, we performed TCR-seq and SARS-CoV-2 antibody assays on 40 participants ten months after their initial vaccine dose (Figure 5A, Data source). Among the participants, 23 tested negative and 17 tested positive for antibodies. The dataset was divided into training and test sets in an 8:2 ratio, and 10 randomized replicate experiments were conducted. In each experiment, the feature selection model was trained using 5-fold cross-validation (Figure 5A, Data distribution). The feature selection process (Figure 5A, Feature selection) involved two steps: (1) an independent t-test was conducted on 1,173,221 CDR3 sequences to identify those with p < 0.05, and (2) the lasso feature selection method was used to further reduce the feature set, minimizing overfitting due to multicollinearity. Logistic regression was chosen as the classification method (Figure 5A, Model), as it demonstrated superior performance on sparse matrices compared to other machine learning approaches such as Random Forest (RF) and support vector machine (SVM)(37). Through ten replicate experiments, 518 antibody-associated CDR3 sequences were identified, including both antibody-positive and antibody-negative sequences. These antibody-positive CDR3 sequences were significantly correlated with antibody-positive participants (Figure 5B), demonstrating the reliability of our in-silico identification process.

The main purpose of this study was to determine whether vaccine antigen-positive CDR3 sequences could serve as biomarkers for detecting antibody responses. Using these identified sequences, we developed a machine learning model to classify participants based on their antibody status (positive or negative). In the 5-fold cross-validation tests, the model achieved a mean AUC of 0.96 between the two groups (Figure 5C). The model also performed exceptionally well in terms of precision, accuracy, F1 score, and recall (Figure 5D).

In this context, the relationship between TCRs and antibodies reveals that TCRs possess binary predictive power with respect to antibody status. While TCRs can indicate whether an individual is antibody-positive or -negative, they do not predict the quantitative level of antibodies. Prior research has also established correlations between T cell responses and antibody test results. Additionally, TCR repertoires are emerging as promising biomarkers for evaluating vaccine efficacy and disease detection. For instance, they have been used to classify cytomegalovirus (CMV)(38) and COVID-19 statuses(39,40). Furthermore, the application of TCR-seq and machine learning has the potential to significantly advance our understanding of immune responses to vaccination and could inform the development of more effective, targeted vaccines for future infectious diseases.

### Characteristics analysis of antibody-associated CDR3 sequences

Subsequently, we examined the features of these antibody-associated CDR3 sequences in samples collected during the T3 stage. Our analysis identified 528 key CDR3 sequences (Figure 6B), encompassing both antibody-positive and antibody-negative sequence. The antibody-positive CDR3 sequences had a higher frequency in antibody-positive samples (RR>1) (RR, relative risk), while the antibody-negative CDR3 sequences were more prevalent in antibody-negative samples (RR<1) (Figure 6A). Notably, 96.9% of the antibody-positive CDR3 sequences were predicted to be SARS-CoV-2-specific according to the DeepAir-Cov19 model. Furthermore, when we mapped these sequences to manually curated TCR specificity databases, we observed that CDR3 sequences with higher publicity were more likely (34%) to be identified in the Immune Epitope Database (IEDB), which catalogs experimentally validated antibody and T cell epitopes across species (Figure 6B). The SARS-CoV-2 genome consists of a single strand of RNA encoding various structural and non-structural proteins(41). Four structure proteins (S, E, M, N) (Figure 6C, green stripes ORF 1ab protein) encapsulate the RNA genome within a lipid bilayer membrane to form infectious viral particles, while sixteen non-structural proteins (Figure 6C, purple stripes) and six accessory proteins (Figure 6C, green stripes) facilitate viral replication and immune evasion(22). The antibody-positive CDR3 sequences identified in the IEDB predominantly targeted ORF 1ab protein (sixteen non-structural proteins), ORF 7b protein (accessory protein), and spike protein (structure protein) (Figure 6D).

**Figure 6.**
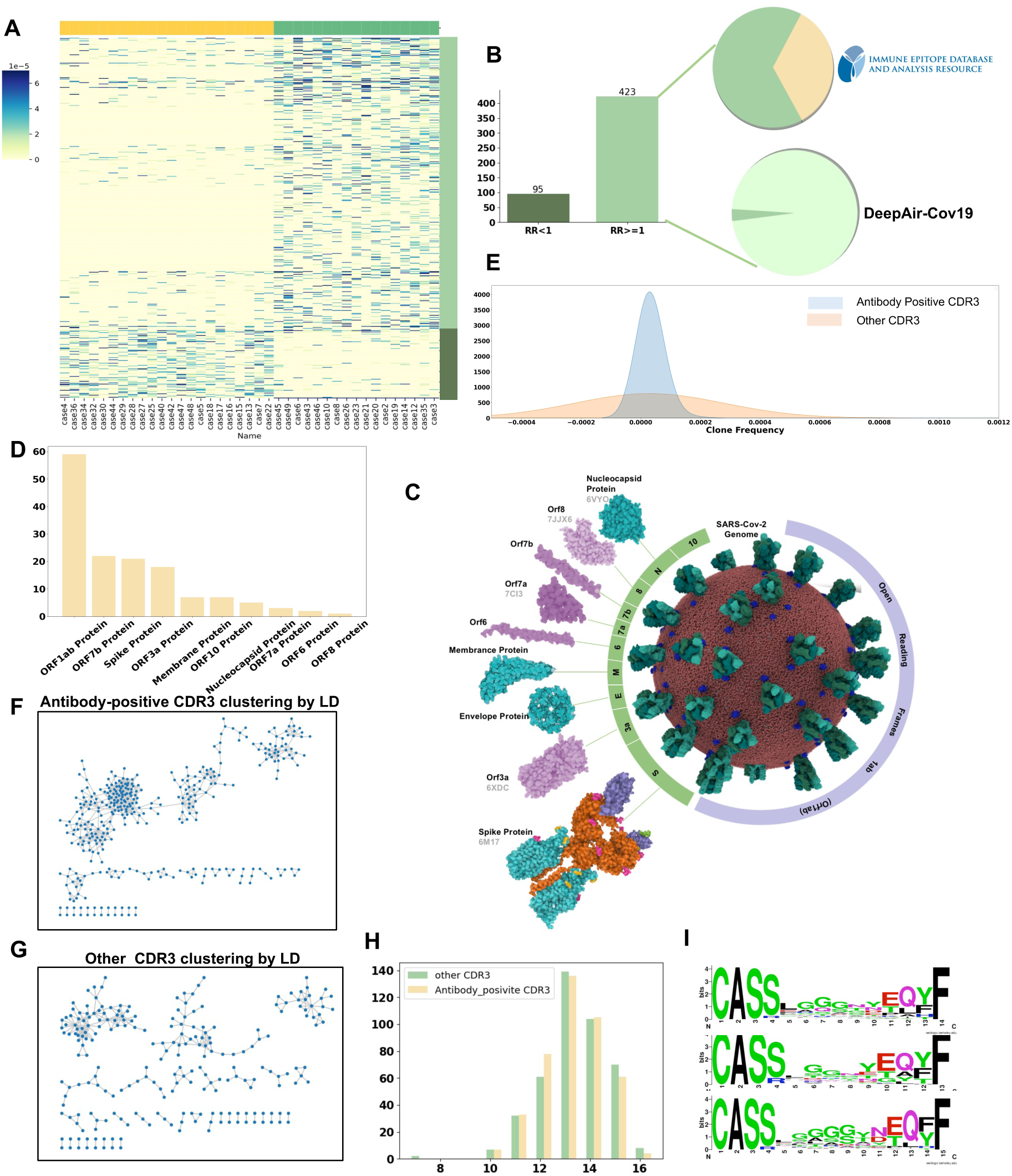
Characteristics analysis of antibody-associated CDR3 sequences of TCRs. (**A**) All antibody-positive TCRβ CDR3 sequences frequency in all participants. Yellow annotation columns are antibody-negative participants. Green annotation columns are antibody-positive participants. Light green annotation lines of the CDR3 group present RR>1. Dark green annotation lines of the CDR3 group present RR<=1. (**B**) Antibody-positive CDR3 sequences are divided into antibody-positive CDR3 sequences (RR>1) and antibody-negative CDR3 sequences (RR<1). 34% of these antibody-positive CDR3 sequences were experimentally validated in IEDB and 96.9% of these antibody-positive CDR3 sequences were predicted to be SARS-CoV-2 specific. (**C**) The genome structure and antigenic proteins of SARS-CoV-2. Purple stripe is open reading frames (ORF) 1ab containing sixteen non-structural proteins. The green stripe contains four structure proteins (S, E, M, N) and six accessory proteins. (**D**) The 10 antigen proteins with antibody-specific CDR3 sequences. (**E**) The comparison results between antibody-positive CDR3 sequences and other control CDR3 sequences by frequency distribution (Kolmogorov-Smirnov test). (**F**) Antibody-positive CDR3 sequences clustering using Levenshtein distance. The dot represents a CDR3, and two CDR3s are clustered together if the Levenshtein distance is less than 3. Dot represents a CDR3 sequence. A line connected the CDR3 sequence in a cluster. (**G**) Antibody-unassociated CDR3 sequences with random sampling clustering using Levenshtein distance. (**H**) CDR3 amino acid length distribution. Antibody-positive CDR3s’ motif with length from 13 to 15 amino acids. (**I**) Antibody-positive CDR3s’ amino acids motif with length from 13 to 15 amino acids

To further investigate the differences between antibody-positive and unrelated CDR3 sequences, we randomly selected unrelated CDR3 sequences from the full dataset, excluding the antibody-associated sequences, ensuring an equal number of unrelated sequences for comparison. Through clone frequency distribution analysis using the Kolmogorov-Smirnov test, we found that antibody-positive CDR3 sequences exhibited significantly higher frequencies and were more clustered than unrelated sequences (Figure 6E). We also explored the sequence similarities within antibody-positive CDR3s using Levenshtein distances, which revealed that these sequences were significantly enriched in large clusters (Figure 6F), a phenomenon not observed in the unrelated CDR3 sequences (Figure 6G). Our data further indicated that the lengths of antibody-specific CDR3 sequences were predominantly between 12 and 15 amino acids, shorter than those of unrelated sequences (Figure 6H). Additionally, glycine was found to play a particularly important role in the diversity of the CDR3 region (Figure 6I). These findings underscore the unique characteristics of antibody-positive CDR3 sequences and suggest that these clones may be critical in the immune response to SARS-CoV-2.

## Discussion

The success of SARS-CoV-2 inactivated vaccines has been remarkable(42,43), yet a deeper understanding of the TCR-mediated immune responses these vaccines elicit remains critical. Such insights could enhance our knowledge of (i) the mechanisms of TCR-based protective immunity, (ii) the longevity of immune memory generated by the vaccines, and (iii) the immunological characteristics of TCR responses that may inform future vaccine designs aimed at optimizing antibody production. In this study, we performed a longitudinal analysis on 42 vaccinated individuals over a span of 10 months following their initial immunization. Our results demonstrate that inactivated vaccines promote the expansion of epitope-specific TCR clones, which significantly enhance SARS-CoV-2-specific immune responses after two vaccine doses.

Accumulating evidence obtained from patients(44–46) and vaccine studies(27,47,48) indicates a central role of T-cell immunity in COVID-19, with T-cell response frequency, intensity, and diversity correlating with SARS-CoV-2 antibody titers and clinical outcomes. Previous studies have documented the generation of spike-specific CD8+ T cells, strong T follicular helper cells, and type 1 helper T cells after vaccination(49), yet few have comprehensively analyzed post-vaccination TCR repertoires. Despite reports of minimal differences in the frequency, phenotype, or TCR motifs between epitope-specific CD8+ T cells generated by natural infection versus vaccination(50), our study extends this understanding by tracking the evolution of TCR repertoires post-vaccination. Using TCR-seq, deep learning models, and machine learning algorithms, we observed that TCR responses persisted beyond the duration of antibody production, suggesting that T cells induced by inactivated vaccines can provide prolonged protection, lasting over ten months. Furthermore, our findings indicate that TCR repertoires can predict antibody neutralization outcomes, underscoring the potential of TCRs as biomarkers for personalized vaccine efficacy assessment.

Measuring T-cell responses to vaccination could provide predictive insights into an individual’s likelihood of developing protective antibody responses, potentially enabling the identification of individuals at higher risk of vaccine failure(51). This predictive capacity could facilitate the development of more targeted vaccination strategies. The growing recognition of T-cell involvement in vaccine-induced immunity reflects their essential role in identifying and eliminating infected cells, thus preventing the spread of infectious diseases. Our study contributes to this expanding body of evidence, showing that T cells not only provide rapid protection but also sustain this protection over time. This is particularly relevant for emerging pathogens, which pose significant global health threats. Effective vaccines for such pathogens often take years to develop, during which time widespread infection and severe disease can occur(52). By leveraging T cells, we may be able to accelerate vaccine development and offer more rapid protection. Moreover, T cells, with their ability to recognize a broad array of pathogens, present an opportunity to develop vaccines that offer wide-ranging protection against multiple infectious diseases(53). This is in contrast to traditional vaccines, which typically target a specific pathogen or strain. By deploying a technique akin to the one used in this study to assess T cells activation, we may be able to create vaccines that provide broader and more durable protection against a variety of infectious diseases.

Developing vaccines for highly variable viruses, like SARS-CoV-2, remains a significant challenge. These viruses mutate rapidly, complicating the development of long-lasting immunity through conventional vaccination approaches(54). Our research presents a novel strategy for addressing this issue by identifying epitope-specific TCRs, enabling the evaluation of vaccine efficacy across viral variants. This approach could pave the way for more effective and long-lasting vaccines, ultimately reducing the disease burden caused by such viruses. Additionally, this method could be applied to other highly variable viruses, such as influenza and HIV, offering a versatile tool for vaccine development and evaluation.

While this study demonstrates the potential advantages of TCR-seq over traditional techniques, it also suffers certain limitations. First, TCR-seq does not directly assess viral clearance or disease resistance. The relationship between TCR responses and vaccine efficacy requires further exploration. Second, the study’s small cohort size limits the generalizability of our findings; larger-scale studies are necessary to validate these results. Finally, TCR sequencing remains more expensive and time-consuming compared to antibody-based techniques like ELISA. These concerns must be considered when interpreting TCR-seq data and evaluating their clinical relevance.

## Conclusions

Our study introduces the DeepAir-Cov19 model, a large language model specifically designed to identify SARS-CoV-2-specific TCRs with high accuracy (AUC = 0.94). Using this model, we revealed significant differences in TCR profiles between antibody-negative and antibody-positive individuals before and after vaccination, demonstrating that pre-vaccination immune status can directly predict vaccine efficacy. Notably, SARS-CoV-2-specific TCRs expanded to peak levels after the second vaccine dose and remained detectable in most subjects for up to 10 months post-vaccination.

Furthermore, we identified three specific V genes, 20 V-J combinations, and three epitopes associated with these responses. By leveraging vaccine-specific TCRs as biomarkers, we developed a predictive model for antibody levels with a mean AUC of 0.96, highlighting the complementary roles of T cells and antibodies in vaccine-mediated protection. These findings provide new insights into the longitudinal dynamics of SARS-CoV-2-specific TCRs and underscore the potential of TCR-based biomarkers for evaluating vaccine potency.

In conclusion, our study emphasizes the critical role of T-cell responses in vaccine efficacy evaluations. Considering both humoral and cellular immunity is essential for a comprehensive assessment of vaccine-induced protection. Further research is needed to fully elucidate the role of T cells in long-term immune memory and to optimize future vaccination strategies.

## Declarations

### Ethics approval and consent to participate

All procedures performed in studies involving human participants were in accordance with the ethical standards of the institutional research committee and with the 1964 Helsinki Declaration and its later amendments or comparable ethical standards. All participants provided written informed consent, and the study was approved by the Ethics Committee of Guangzhou Laboratory Medical Research and Lunar Review 2021, No. 78.

### Consent for publication

Written informed consent for publication was obtained from each participant.

### Availability of data and materials

All the data analyzed in this study are available within the paper and supplementary files. The TCR sequence data for all the samples used in this study are available at Google Drive (https://drive.google.com/file/d/176Oq7yrvQz-x53meF0C7PqL4ILc7I8NF/view?usp=sharing).

### Authors’ contributions

ZHW conceived the project and directed the study with input from all authors. ZHW, BH, WMZ, WHL and JHYwere involved in study design. BH provided study oversight. LPC, XX and JB provided data access. ZHW and YZ conducted data analysis. ZHW and BH wrote the manuscript. XX, YLS, SX, CCQ, JB, ZPZ were involved in data interpretation and editing the manuscript.

## Acknowledgements

Not applicable.

## Supplementary Information

Additional file 1. Supplementary Figures: Figs. S1-S4.

Additional file 2. Supplementary Tables

